# Fibroblastic reticular cells predict response to immune checkpoint inhibitors

**DOI:** 10.1101/2020.02.19.955666

**Authors:** Daniele Biasci, James Thaventhiran, Simon Tavaré

## Abstract

While the role of CD8+ T cells in mediating response to cancer immunotherapy is well established, the role of B cells remains more controversial (1–3). By conducting a large gene expression study of response to immune checkpoint inhibitors (ICI), we show that pre-treatment expression of B cell genes is associated with ICI response independently of CD8+ T cells. However, we discovered that such association can be completely explained by a single gene (*FDCSP*) expressed outside of the B cell compartment, in fibroblastic reticular cells (FRCs), which form the reticular network that facilitates interactions between B cells, T cells and cognate antigens (4–6) and are required to initiate efficient adaptive immune responses in secondary lymphoid organs (SLO) and tertiary lymphoid structures (TLS) (4, 7). We validated this finding in three independent cohorts of patients treated with ICI in melanoma and renal cell carcinoma. Taken together, these results suggest that *FDCSP* is an independent predictor of ICI response, thus opening new avenues to explain the mechanisms of resistance to cancer immunotherapy.

---

Recently, it has been reported that expression of B cell specific genes inside tumours is associated with response to immune checkpoint inhibitors (ICI), thus suggesting an unappreciated biological role for B cells in promoting ICI response (8). However, association between B cell gene expression and ICI response was not statistically significant when other immune populations, including CD8+ T cells, were taken into account (8). For this reason, it remains unclear whether B cells are associated with ICI response independently of CD8+ T cells. Clarifying this point is crucial for the biological interpretation of these results. In fact, expression of B cell specific genes in the tumour microenvironment (TME) is often strongly correlated with markers of CD8+ T cells (9), which express the actual molecular targets of ICI treatment and are a validated predictor of immunotherapy response (10–12). Two additional recent studies on this topic did not address the issue, because they described predictive signatures containing both B cell and CD8+ T cell markers, as well as genes expressed in other immune cell types (13, 14). In order to address this problem, we analysed the entire transcriptome of a larger number of melanoma samples compared to previous studies (*n* = 366), and we then considered only samples collected before commencing treatment with immune checkpoint inhibitors. Moreover, we excluded tumour samples obtained from lymph node metastasis, in order to minimise the risk of capturing unrelated immune populations from the adjacent lymphoid tissue (see supplementary methods). We calculated the correlation between expression of each gene at baseline and subsequent treatment response, as defined in the original studies according to the RECIST criteria (15). A distinct group of genes showed significant association with treatment response, namely: *CR2* (CD21), *MS4A1* (CD20), *CD19, FCER2* (CD23), *PAX5, BANK1, VPREB3, TCL1A, CLEC17A* and *FDCSP* (Fig. 1A and Table S1). Literature reports that these genes are predominantly expressed in B cells (16), with the exception of *FDCSP*, which was found to be expressed follicular dendritic cells (FDCs) isolated from secondary lymphoid organs, but not in B cells (17, 18). In order to identify the cell populations expressing these genes in tumours, we assessed their expression in single-cell RNA sequencing data (scRNAseq) from human cancer samples (19). First, we confirmed that genes identified by our association study were specifically expressed in tumour-associated B cells (Fig. S7). Second, we observed that *FDCSP* was not expressed in tumour-associated B cells (Fig. S1), but identified a subset of fibroblastic cells (*FAP*+ *COL1A1*+ *CD31*-; Fig. S2) expressing markers consistent with FRC identity (*CCL19*+ *CCL21*+ *BAFF*+; 4, 5; Figures 2 and S3 and tables S5 and S6). Finally, we confirmed these results in an independent set of melanoma samples profiled at the single-cell level (20; Figures 2, S4 to S6 and S8). We then sought to determine whether expression of these genes was associated with ICI response independently of CD8+ T cell infiltration. To this aim, we repeated the association analysis after taking into account expression markers of T cells, NK cells, cross-presenting dendritic cells (21) and interferon-gamma response (22; Fig. 1D and table S2). We found that only four genes were associated with ICI response independently of all immune markers considered: *MS4A1* (prototypical B cell marker), *CLEC17A* (expressed in dividing B cells in germinal centers; 23), *CR2* (expressed in mature B cells and FDCs; 24) and *FDCSP* (expressed in FRCs but not in B cells). This result suggested that presence of B cells and FRCs in the tumour microenvironment (TME) before treatment was associated with subsequent ICI response independently of CD8+ T cells (Fig. 1D and Table S7). Remarkably, we observed that *FDCSP* remained significantly associated with ICI response even after B cell markers were taken into account (Fig. 1D) and that no other gene remained significant after *FDCSP* expression was included in the model (Fig. 1D). Taken together, these results suggest that expression of *FDCSP* might be an independent predictor of ICI response. We tested this hypothesis in a completely independent cohort of melanoma patients treated with anti-PD1, either alone or in combination with anti-CTLA4 (25). In this validation cohort, patients with higher expression of *FDCSP* at baseline experienced significantly longer overall survival (log-rank *p* < 0.0001) and progression-free survival (log-rank *p* < 0.0001) after commencing treatment with ICI (Figures 1B and 1C). Moreover, we observed a significant association between expression of *FDCSP* at baseline and subsequent RECIST response (Cochran-Armitage test *p* < 0.0001; Fig. 1E). We sought to replicate this observation in two additional independent cohorts of patients treated with ICI in melanoma (8) and clear cell renal cell carcinoma (26) and found similar results (Figures 1F and 1G). On the other hand, testing B cell markers in the validation cohorts produced less consistent results (Fig. S9). Finally, we performed multiple linear regression analysis and found that the association between *FDCSP* and ICI response was independent of CD8+ T cells, CD4+ T cells, B cells, NK cells, myeloid Dendritic cells (mDCs), Macrophages and Plasma cells (Tables 1 and S3). While CD8+ T cells express the molecular targets of ICI, and their presence in the TME is a validated predictor of ICI response (10–12), the role of B cells remains more controversial (1–3). Because T and B cells often co-occur in the TME (27), gene expression studies consistently reported strong correlation between T cells, B cells and other immune cell types, especially in melanoma (9). As a consequence, assessing whether B cells are associated with ICI response independently of CD8+ T cells has been proven challenging (8). By analysing a larger number of samples compared to previous studies, we were able to demonstrate that B cells predict ICI response independently of CD8+ T cells (Fig. 1D and Table S4). This supports the idea that B cells might actively promote immunotherapy response (3), rather than being irrelevant or detrimental to it (1, 2). However, our data also shows that such positive association can be completely explained by a single gene expressed outside of the B cell compartment. We have demonstrated that *FDCSP* is transcribed in a subset of fibroblastic cells up-regulating *CCL19, CCL21* and *BAFF* and down-regulating genes associated with canonical fibroblasts, such as extracellular matrix genes (Fig. 2, Table S5, Table S6), an expression pattern consistent with FRC identity (4, 5). One way to reconcile these observations is to consider that FRCs are required to initiate efficient B and T cell responses (4). In both secondary lymphoid organs (SLO) and tertiary lymphoid structures (TLS), FRCs form reticular networks that facilitate interactions between B cells, T cells and their cognate antigens (5, 6). Homing of immune cells into the network requires interaction between chemokines produced by FRCs (CCL19 and CCL21) and their receptor (CCR7) expressed on T cells, B cells and dendritic cells (28, 29; see also Table S1). Accordingly, depleting FRCs causes loss of T cells, B cells and dendritic cells in both SLO and TLS and decreases the magnitude of B and T cell responses to subsequent viral infection (4, 7). Perhaps more importantly, differentiation of fibroblasts into FRCs occurs early during TLS formation, precedes B and T cell infiltration and can still be observed in *Rag2-/-* mice (7). In this context, our finding that a gene expressed in FRCs is associated with ICI response independently of B and T cell infiltration is coherent with the current understanding of TLS neogenesis. Taken together, these observations suggest that the reported association between B cells and ICI response (8), albeit independent of CD8+ T cells, might ultimately be secondary to the presence of FRCs in the TME. The identification of *FDCSP* as a single marker of ICI response should enable even larger studies to test this conclusion. To our knowledge, this is the first time that FRCs are directly implicated in ICI response, thus opening new avenues to explain the mechanisms of resistance to ICI.

**Fig. 1.**
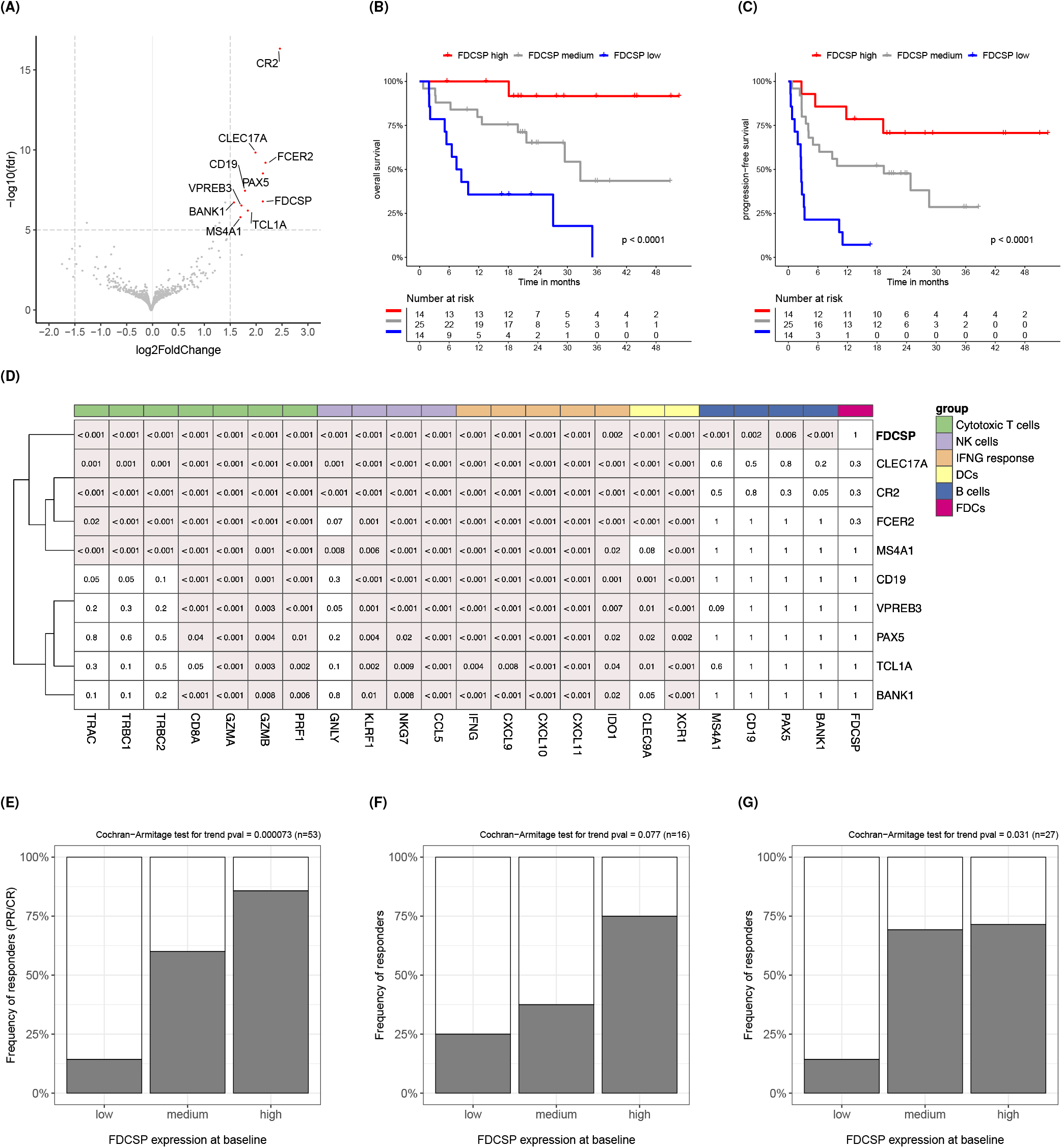
Expression of FDCSP is an independent predictor of response to immune checkpoint inhibitors. **A.** Volcano plot showing genes significantly associated with response to ICI. **B-C** Kaplan-Meier curves showing different overall survival and progression-free survival in melanoma patients after commencing treatment with ICI. Patients from the first validation cohort (25; see supplementary methods) were stratified in three groups according to pre-treatment expression of *FDCSP*. **D** Heatmap showing association between pre-treatment expression of B cell or FRC genes (rows) and subsequent ICI response after the effect of other immune markers (columns) was taken into account. The number inside each cell represents the Wald test p-value for the underlying effect size estimate obtained using the negative binomial model for RNAseq counts implemented in DESEq2 (see supplementary methods). Nominal p-values were corrected for multiple testing using Benjamini–Yekutieli false discovery rate (FDR) and are also reported in Table S7. A red background highlights effect sizes estimated to be greater than zero with FDR < 0.05. In this study, an effect size greater than zero represents a positive association between gene expression and probability of ICI response. **E-G** Association between pre-treatment expression of *FDCSP* and subsequent ICI response in the first (25), second (8) and third validation cohort (26). See supplementary methods.

**Fig. 2.**
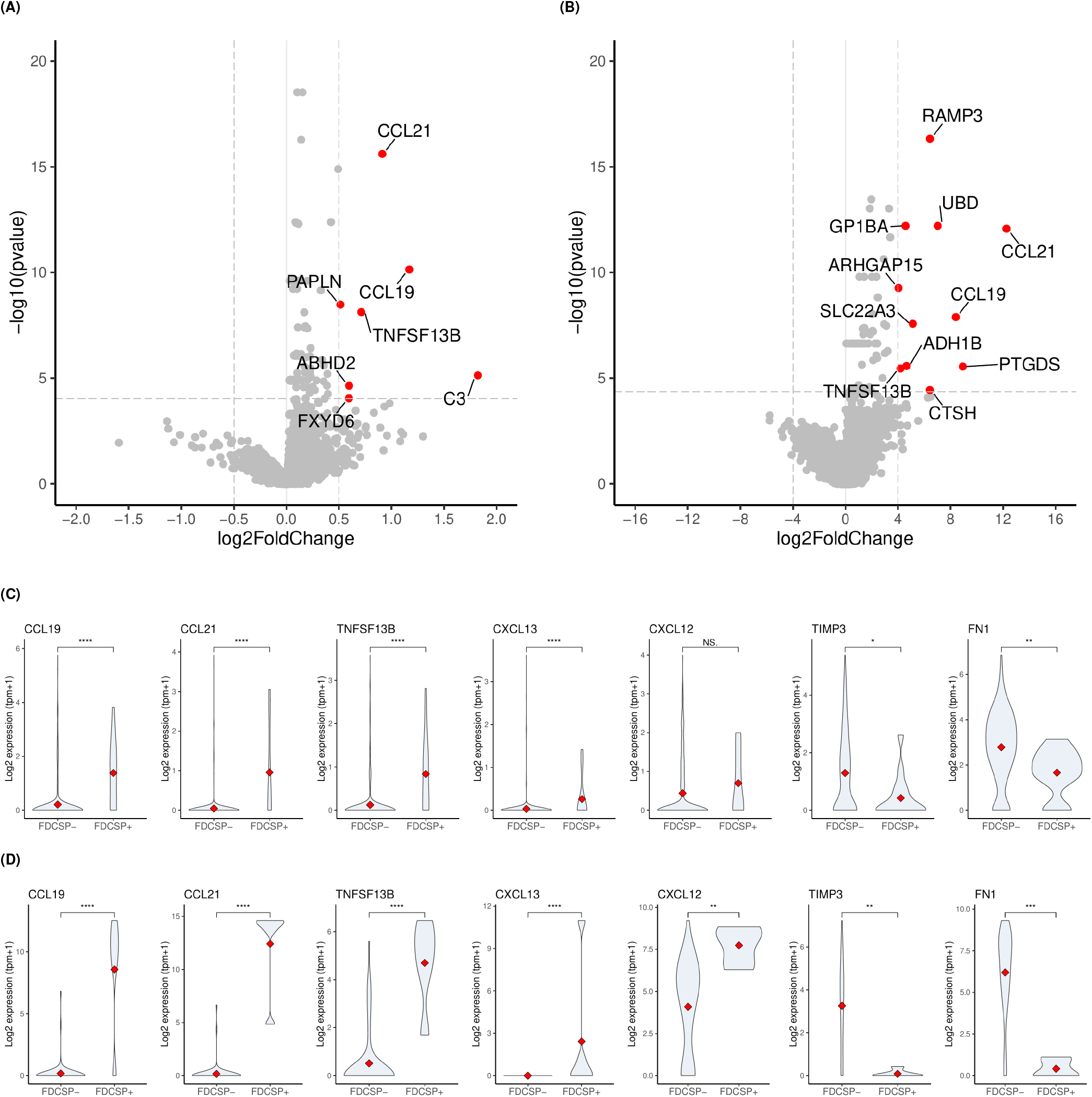
*FDCSP*+ fibroblasts upregulate FRCs markers in lung cancer and melanoma. Differential gene expression analysis comparing single-cell transcriptomes of *FDCSP*+ fibroblasts versus *FDCSP*- fibroblasts obtained from lung cancer and melanoma tumours. The analysis was performed using gene expression data published in (19, 20); see also supplementary methods, Table S5 and Table S6. **A.** Volcano plot representing genes differentially expressed between *FDCSP*+ and *FDCSP*- fibroblasts shows that *FDCSP*+ fibroblasts significantly upregulate expression of FRCs markers (*CCL19, CCL21, TNFSF13B/BAFF)* in lung cancer. Genes considered significantly up-regulated (log_2_ (*fc*) > max(log_2_ (*fc*))/4 and FDR < 0.05) are marked in red; dotted lines represent chosen log_2_ (*fc*) and FDR cutoffs. **B.** Volcano plot representing genes differentially expressed between *FDCSP*+ and *FDCSP*- fibroblasts and confirming significant up-regulation of FRCs markers in melanoma. **C.** Violin plots comparing distribution of gene expression values in *FDCSP*+ and *FDCSP*- fibroblasts in lung cancer. The plots shows that *FDCSP*+ fibroblasts significantly upregulate cytokines relevant to FRCs function while down-regulating genes involved in secretion of the extracellular matrix (*TIMP3, FN1*). Statistical significance was assessed using unpaired two-samples Wilcoxon test. A red diamond represents the median value of each distribution. **D.** Comparing *FDCSP*+ and *FDCSP*- obtained from melanoma tumours leads to similar results (see panel C).

**Table 1.** Multiple linear regression analysis of ICI response. Expression values of cell type specific markers were summarised by calculating their geometric mean. The resulting summarised values were used as independent variables in a multiple linear regression model aimed at explaining ICI response. The table reports the regression coefficient estimates obtained by fitting the model to all pre-treatment melanoma samples available for this study (see Supplementary methods). The probability of observing estimates different from zero merely by chance was calculated by leveraging on the null distribution of the t-test statistic (*Pr*(> |*t*|)). The results show that expression of *FDCSP* is significantly associated with ICI response independently of genes expressed in major immune cell types. Asterisks indicate the level of statistical significance: *** indicates p-value ≤ 0.001, ** indicates p-value ≤ 0.01, * indicates p-value ≤ 0.05.

|  | Cell type | Markers | Estimate | Std. Error | t value | Pr(>\|t\|) |  |
| --- | --- | --- | --- | --- | --- | --- | --- |
| 1 | CD8+ T cells | CD8A, CD8B, TRAC | 0.032 | 0.078 | 0.40 | 0.69 |  |
| 2 | CD4+ T cells | CD4, TRAC | 0.057 | 0.110 | 0.52 | 0.60 |  |
| 3 | NK cells | GNLY, KLRF1 | -0.089 | 0.092 | -0.96 | 0.34 |  |
| 4 | B cells | MS4A1, CD19 | 0.044 | 0.049 | 0.90 | 0.37 |  |
| 5 | Macrophages | CD163, MARCO | -0.014 | 0.049 | -0.29 | 0.77 |  |
| 6 | Dendritic cells | CD1A, CD1B, CD1E, LILRA4 | -0.053 | 0.075 | -0.70 | 0.48 |  |
| 7 | FDCSP+ cells | FDCSP | 0.071 | 0.027 | 2.60 | 0.0092 | ** |

## Appendix S1: supplementary methods

### Bulk RNAseq: discovery cohorts

The RNAseq datasets used in this study as discovery cohorts were downloaded from the following published sources: data for 27 melanoma samples from patients that went on to receive ICI treatment was downloaded from the NIH Genomics Data Common Portal, study TCGA-SKCM (1, 2); data for 42 pre-ICI melanoma samples was downloaded from dbGaP (3), study phs000452.v2.p1 (4); data for 109 melanoma samples collected before or during ICI treatment was downloaded from Gene Expression Omnibus (GEO;5), series GSE91061 (6); data for 28 melanoma samples collected before ICI treatment was downloaded from GEO, series GSE78220 (7); data for 37 melanoma samples collected before or during ICI treatment was downloaded from GEO, series GSE115821 (8). Clinical annotations for each sample (i.e. type of ICI treatment received, sample collection time point, tumour site, treatment response) were obtained from the original publications. Samples collected after commencing treatment with ICI, samples obtained from lymph node metastasis, and samples for which treatment response status was not available were excluded from subsequent analyses, leaving 137 pre-treatment samples available for this study. This manuscript reanalyses data from already published studies: ethics and consent information can be found in the original studies.

### Bulk RNAseq: validation cohorts

The RNAseq datasets used in this study as validation cohorts were downloaded from the following published sources. *First validation cohort:* data for 91 melanoma samples collected before or during ICI treatment was downloaded from the European Nucleotide Archive (ENA;9), study PRJEB23709 (10). *Second validation cohort:* data for 16 melanoma samples collected before ICI treatment was obtained from (11). *Third validation cohort:* data for 33 clear cell renal cell carcinoma samples collected before ICI treatment was obtained from (12). Clinical annotations for each sample were obtained from the original publications. Samples collected after commencing treatment with ICI and samples for which treatment response status was not available were excluded from subsequent analyses.

### Bulk RNAseq: gene expression quantification

Sequences of all known human transcripts were downloaded from Ensembl release 97 (13). Transcript quantification was performed using Kallisto ver. 0.43 (14)and gene-level count matrices for use in DESeq2 (15) were calculated using tximport (16) as recommended by DESeq2 authors (17). All subsequent analyses on gene expression were performed using R 3.5.0 (18). For differential expression analysis, raw counts were used directly in DESeq2 as recommended (17). For other downstream analyses (i.e. survival curves, multiple correlation analysis and others) counts data were transformed using the variance stabilizing transformation (19) as implemented in DESEq2 (17) and potential systematic differences between cohorts were corrected using ComBat (20) as implemented in the sva package (21). The methods described in this section were used to quantify gene expression in all datasets used in this study as discovery cohorts and for the first validation cohort, for which original FASTQ files were available. For the second and third validation cohort (10), we used the gene expression values provided in the original publications (11, 12).

### Bulk RNAseq: genes associated with ICI response

In the discovery cohorts, association between gene expression and ICI response was calculated using the DESeq2 model based on the negative binomial distribution (15). Raw counts for each gene were used directly in the model as recommended by DESeq2 authors (17). We used the ICI response status for each patient as reported in the original study (1, 4, 6–8) according to the RECIST criteria (22). RECIST response categories were encoded as a numerical score as follows: PD (progressive disease) = −1, SD (stable disease) = 0, PR (partial response) = 0.5, CR (complete response) = 1. When we assessed the use of a different numerical score (i.e. PD= 0, SD=0.33, PR= 0.66, CR= 1), we obtained similar results. In order to account for systematic differences between cohorts, a categorical variable encoding the cohort of origin for each sample was included as a covariate in the DESeq2 association model. In order to take into account the effect of known immune markers (Fig. 1D), the association analysis was repeated after their expression level was calculated according to DESeq2 documentation (17) and added to the model as an additional covariate.

### Bulk RNAseq: survival curves

Survival curves were plotted using the R package survminer (23). Patients were stratified in three groups according to the baseline expression of the gene of interest: the high expression group was defined as containing the 25% of samples with highest expression, the low expression group was defined as containing the 25% of samples with lowest expression, and the medium expression group was defined as containing the remaining samples. Statistical significance of the observed difference between groups was assessed using the Logrank test (24).

### Single-cell RNAseq: original data sets

Expression values (normalised counts) for 54773 single cells isolated from non-small-cell lung cancer (NSCLC) tumour samples were downloaded from GEO, series GSE127465 (25). Cell type annotation and a two-dimensional visualisation (SPRING plot, 26) of single-cell transcriptomes were also obtained from the original publication (25) and used for subsequent analyses. Expression values (transcripts per million) for 7186 single cells isolated from melanoma samples were downloaded from GEO, series GSE115978 (27). Cell type annotation and a two-dimensional visualisation (tSNE plot,28) of single-cell transcriptomes were obtained from the original publication (27) and used for subsequent analyses.

### Single-cell RNAseq: differentially expressed genes

Expression values obtained from GSE127465 and GSE115978 were transformed in logarithmic scale (*y* = log_2_(*x* + 1)). Cells annotated as fibroblasts in the original study were selected and used to compare fibroblasts expressing *FDCSP* (*FDCSP*+) against fibroblasts not expressing *FDCSP* (*FDCSP*-). Statistical significance for gene expression differences observed between these two groups was assessed using unpaired two-samples Wilcoxon test (29). Genes were ranked using a numerical score which takes into account both p-value and log_2_(*fc*): 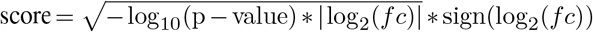. Results for GSE127465 (non-small cell lung cancer) and GSE115978 (melanoma) were meta-analysed using the RankProduct method (30, 31) and are presented in Tables S5 and S6.

### Code availability

The authors declare that the code used for this study is available upon request.

## Appendix S2: supplementary figures

**Fig. S1.**
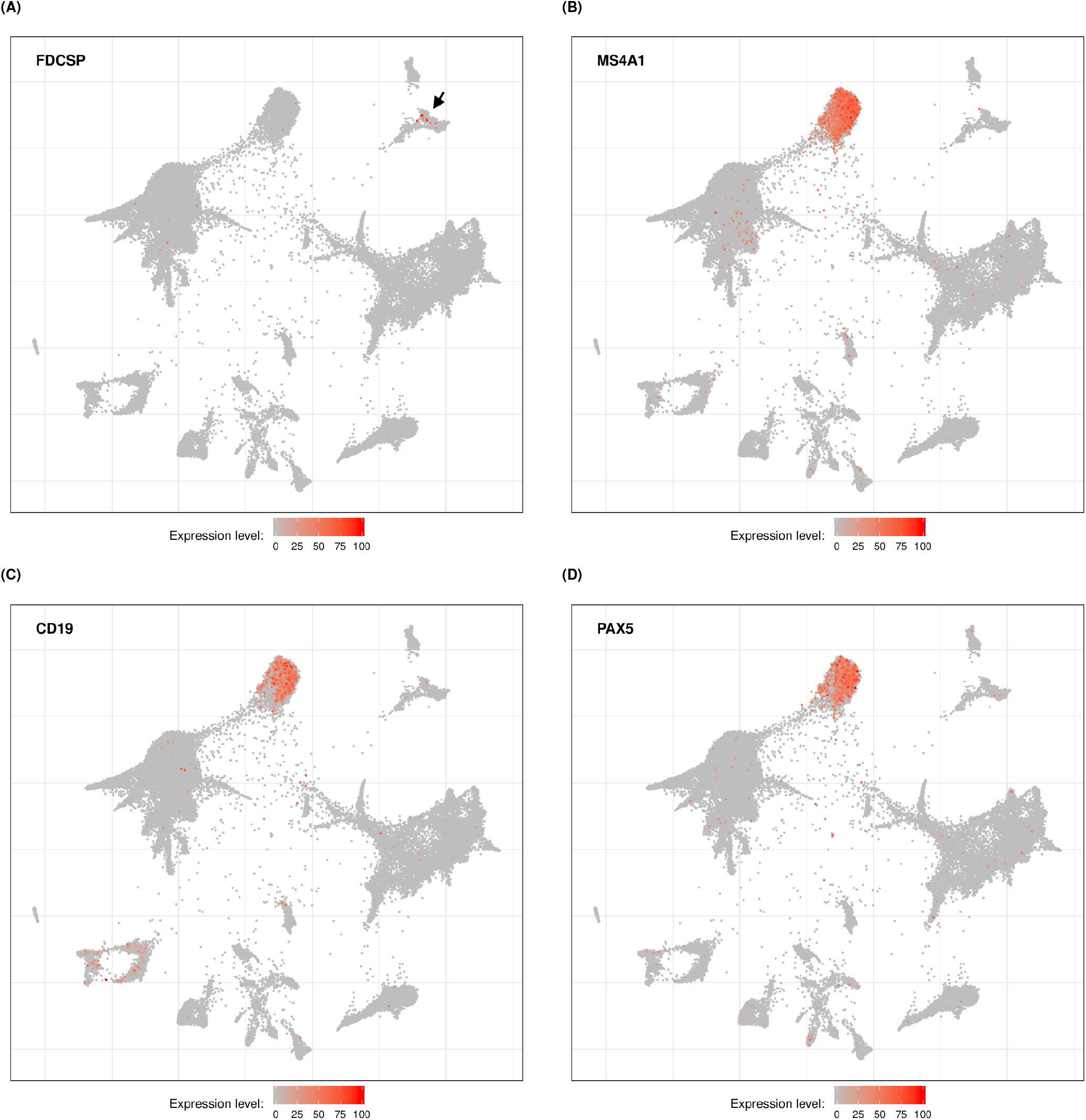
FDCSP is not expressed in B cells infiltrating lung cancer tumours. Two-dimensional visualizations (SPRING plots) of single-cell RNA sequencing data obtained from lung cancer tumours and published in (25). Each dot represents the transcriptional profile of a single cell. Cells closely associated in each plot are more likely to transcribe similar genes and might thus belong to the same cell type. Expression values and coordinates for each dot were obtained from the original study. The color scale indicates expression level for a particular gene ranging from 0 (minimum expression value observed in the dataset for that gene, usually corresponding to non detectable expression) to 100 (maximum expression value observed in the dataset for that gene). Expression of *FDCSP* is observed in a group of cells indicated by the black arrow in (**A**) well distinct from cells expressing prototypical B cell markers (**B-D**).

**Fig. S2.**
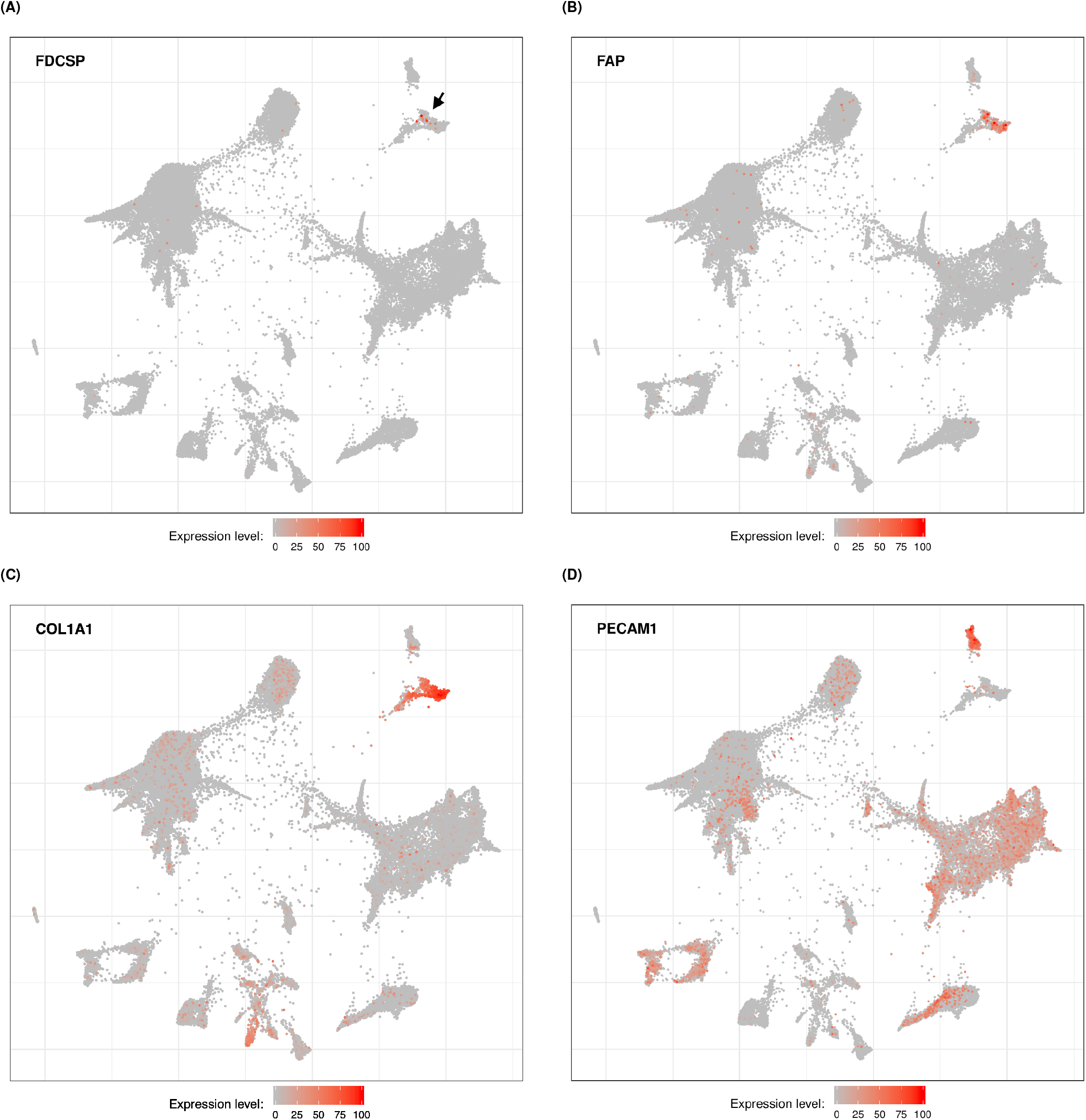
FDCSP is expressed in a subset of fibroblasts in lung cancer tumours. Two-dimensional visualizations (SPRING plots) of single-cell RNA sequencing data obtained from lung cancer tumours and published in (25). Each dot represents the transcriptional profile of a single cell. Cells closely associated in each plot are more likely to transcribe similar genes and might thus belong to the same cell type. Expression values and coordinates for each dot were obtained from the original study. The color scale indicates expression level for a particular gene ranging from 0 (minimum expression value observed in the dataset for that gene, usually corresponding to non detectable expression) to 100 (maximum expression value observed in the dataset for that gene). **A.** Expression of *FDCSP*. The black arrow indicates a group of transcriptionally related cells containing *FDCSP+* cells. **B.** Expression of *FAP*. **C.** Expression of *COL1A1*. **D.** Expression of *PECAM1/CD31*.

**Fig. S3.**
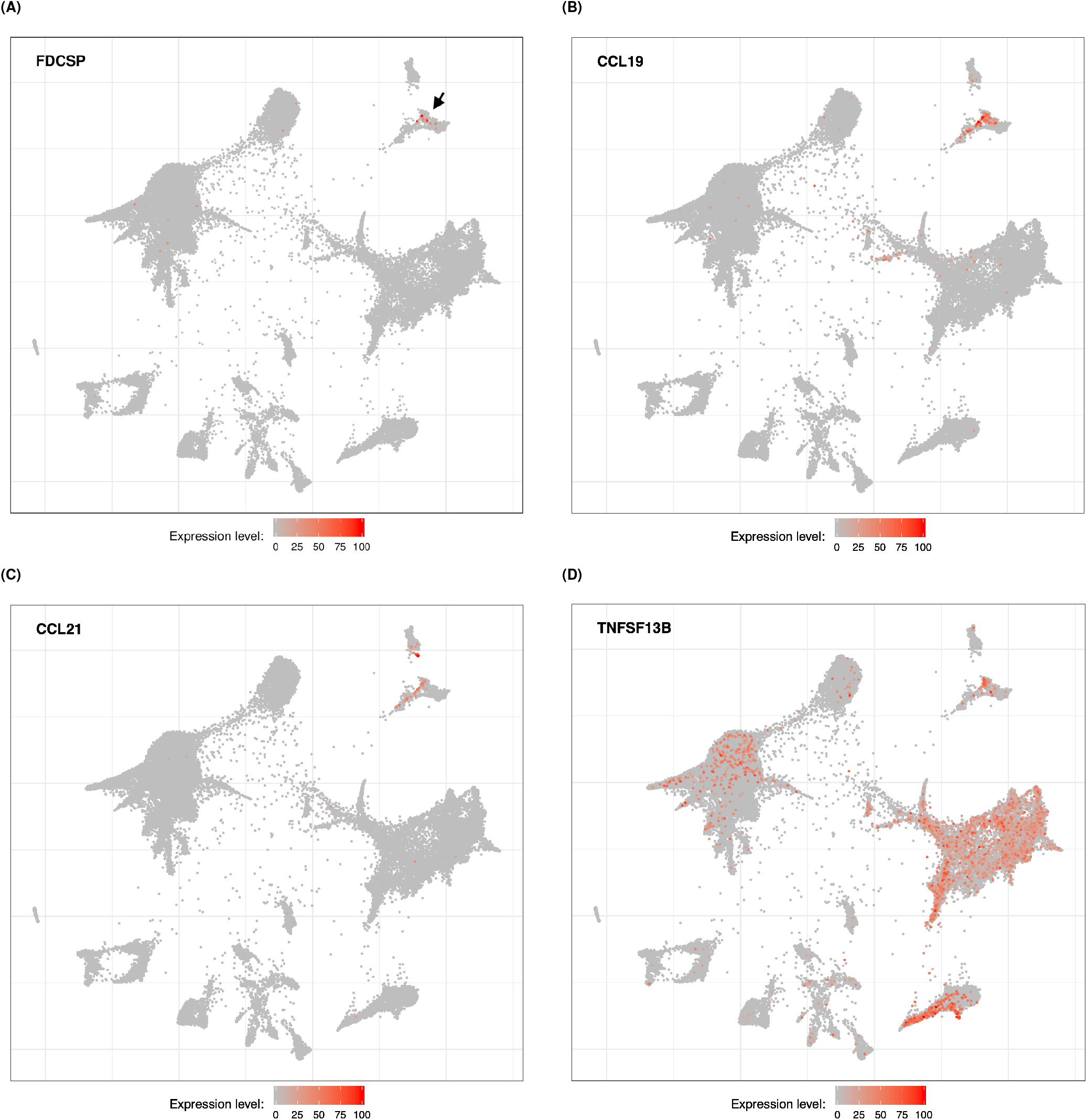
FDCSP+ cells express markers of FRCs in lung cancer tumours. Two-dimensional visualizations (SPRING plots) of single-cell RNA sequencing data obtained from lung cancer tumours and published in (25). Each dot represents the transcriptional profile of a single cell. Cells closely associated in each plot are more likely to transcribe similar genes and might thus belong to the same cell type. Expression values and coordinates for each dot were obtained from the original study. The color scale indicates expression level for a particular gene ranging from 0 (minimum expression value observed in the dataset for that gene, usually corresponding to non detectable expression) to 100 (maximum expression value observed in the dataset for that gene). **A.** Expression of *FDCSP*. The black arrow indicates a group of transcriptionally related cells containing *FDCSP*+ cells. **B.** Expression of *CCL19*. **C.** Expression of *CCL21*. **D.** Expression of *TNFSF13B/BAFF*.

**Fig. S4.**
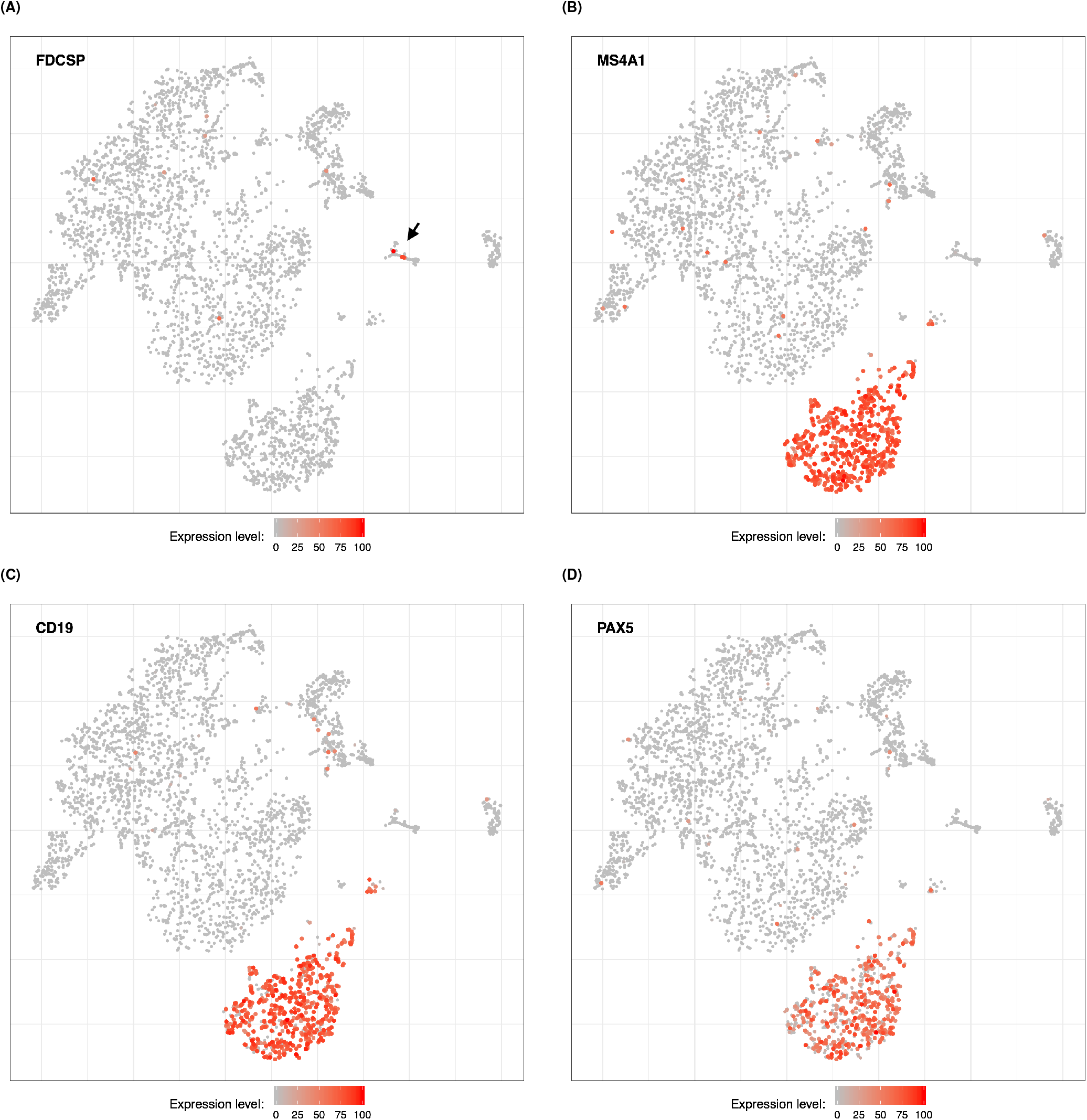
FDCSP is not expressed in B cells infiltrating melanoma. Two-dimensional visualizations (tSNE plots) of single-cell RNA sequencing data obtained from melanoma tumours and published in (27). Each dot represents the transcriptional profile of a single cell. Cells closely associated in each plot are more likely to transcribe similar genes and might thus belong to the same cell type. Expression values and coordinates for each dot were obtained from the original study. The color scale indicates expression level for a particular gene ranging from 0 (minimum expression value observed in the dataset for that gene, usually corresponding to non detectable expression) to 100 (maximum expression value observed in the dataset for that gene). Expression of *FDCSP* is observed in a group of cells indicated by a black arrow in (**A**) well distinct from cells expressing prototypical B cell markers (**B-D**).

**Fig. S5.**
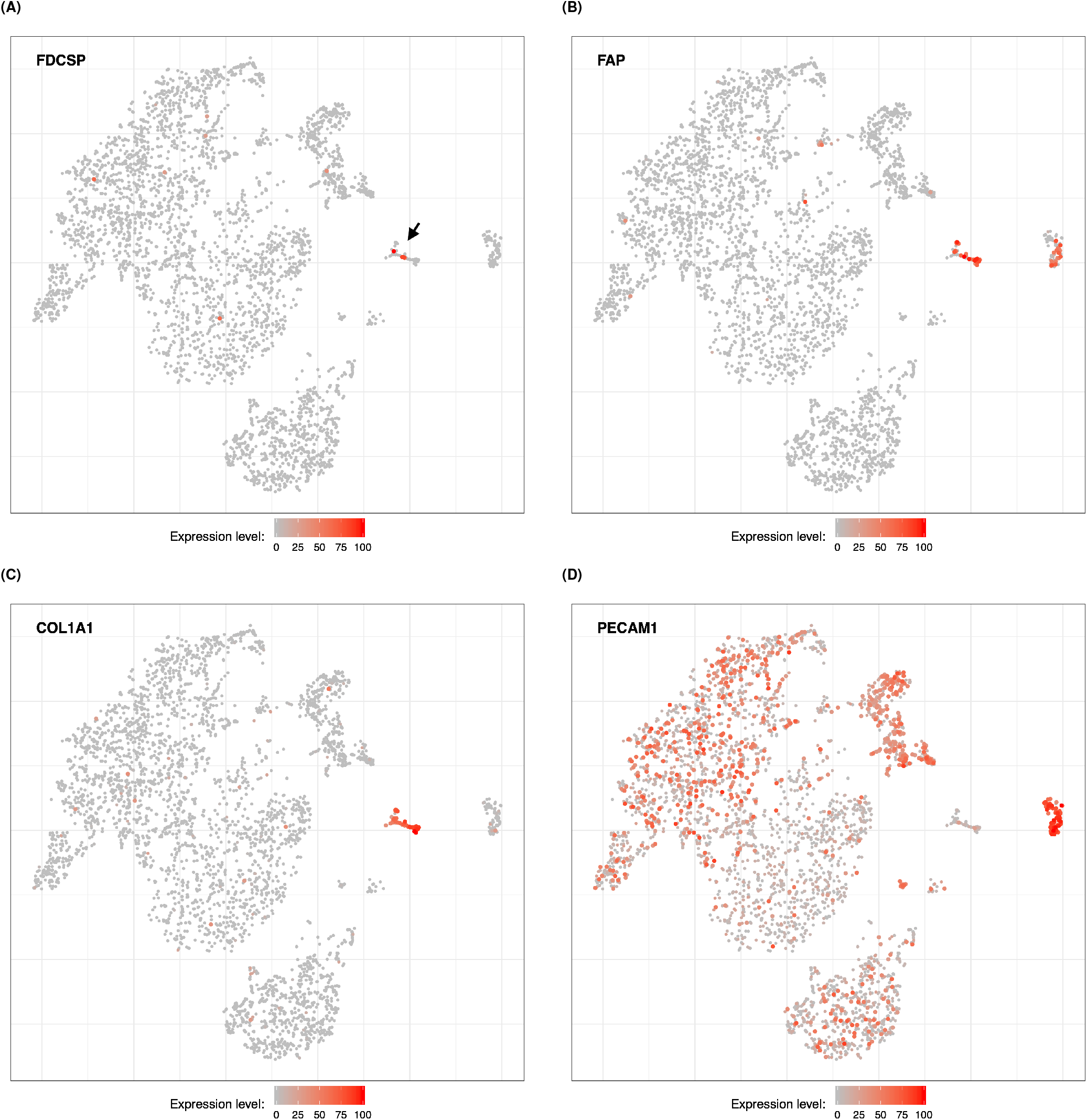
FDCSP is expressed in a subset of fibroblasts in melanoma. Two-dimensional visualizations (tSNE plots) of single-cell RNA sequencing data obtained from melanoma tumours and published in (27). Each dot represents the transcriptional profile of a single cell. Cells closely associated in each plot are more likely to transcribe similar genes and might thus belong to the same cell type. Expression values and coordinates for each dot were obtained from the original study. The color scale indicates expression level for a particular gene ranging from 0 (minimum expression value observed in the dataset for that gene, usually corresponding to non detectable expression) to 100 (maximum expression value observed in the dataset for that gene). **A.** Expression of *FDCSP*. The black arrow indicates a group of transcriptionally related cells containing *FDCSP+* cells. **B.** Expression of *FAP*. **C.** Expression of *COL1A1*. **D.** Expression of *PECAM1/CD31*.

**Fig. S6.**
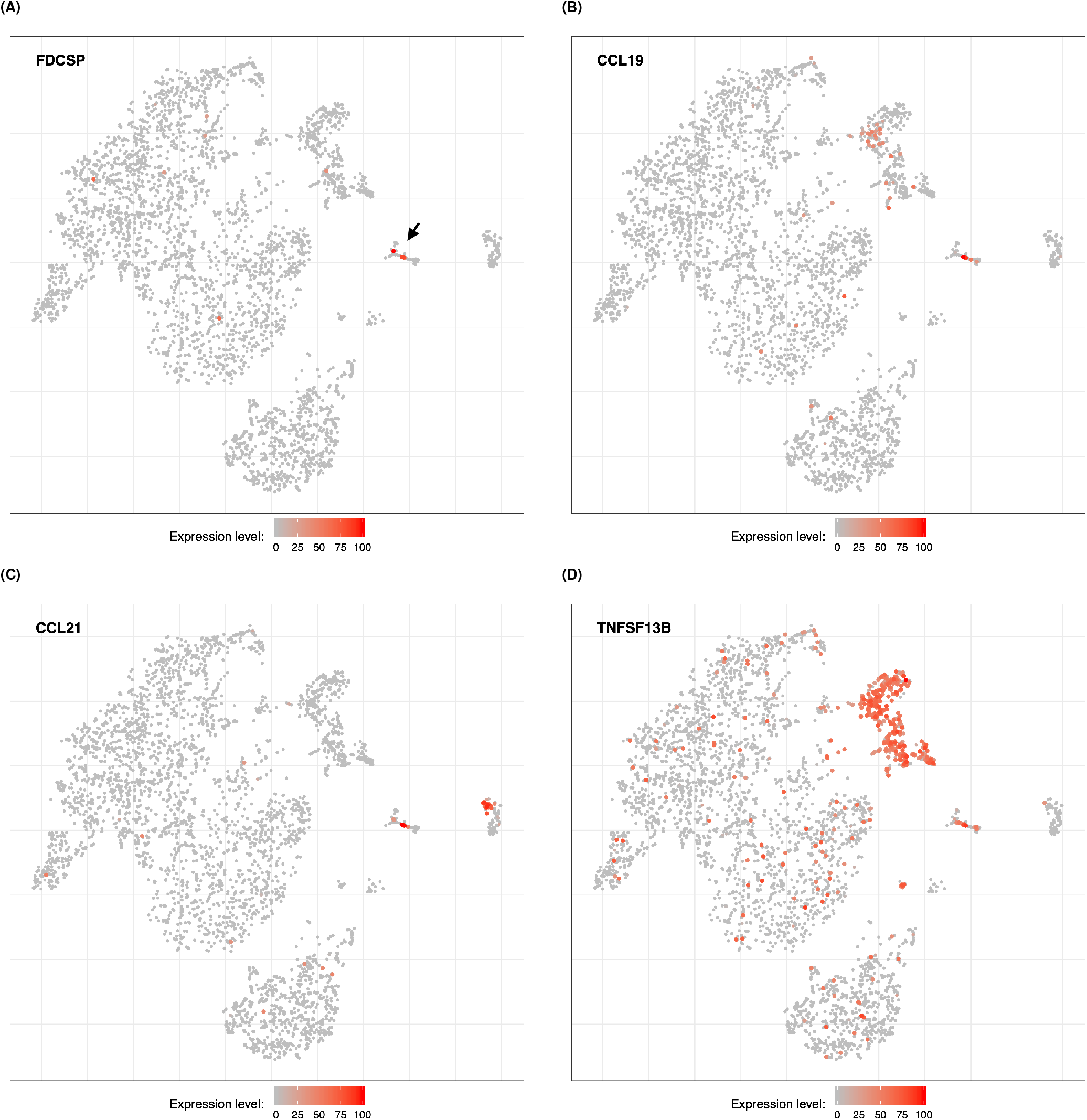
FDCSP+ cells express markers of FRCs in melanoma. Two-dimensional visualizations (tSNE plots) of single-cell RNA sequencing data obtained from melanoma tumours and published in (27). Each dot represents the transcriptional profile of a single cell. Cells closely associated in each plot are more likely to transcribe similar genes and might thus belong to the same cell type. Expression values and coordinates for each dot were obtained from the original study. The color scale indicates expression level for a particular gene ranging from 0 (minimum expression value observed in the dataset for that gene, usually corresponding to non detectable expression) to 100 (maximum expression value observed in the dataset for that gene). **A.** Expression of *FDCSP*. The black arrow indicates a group of transcriptionally related cells containing *FDCSP*+ cells. **B.** Expression of *CCL19*. **C.** Expression of *CCL21*. **D.** Expression of *TNFSF13B/BAFF*.

**Fig. S7.**
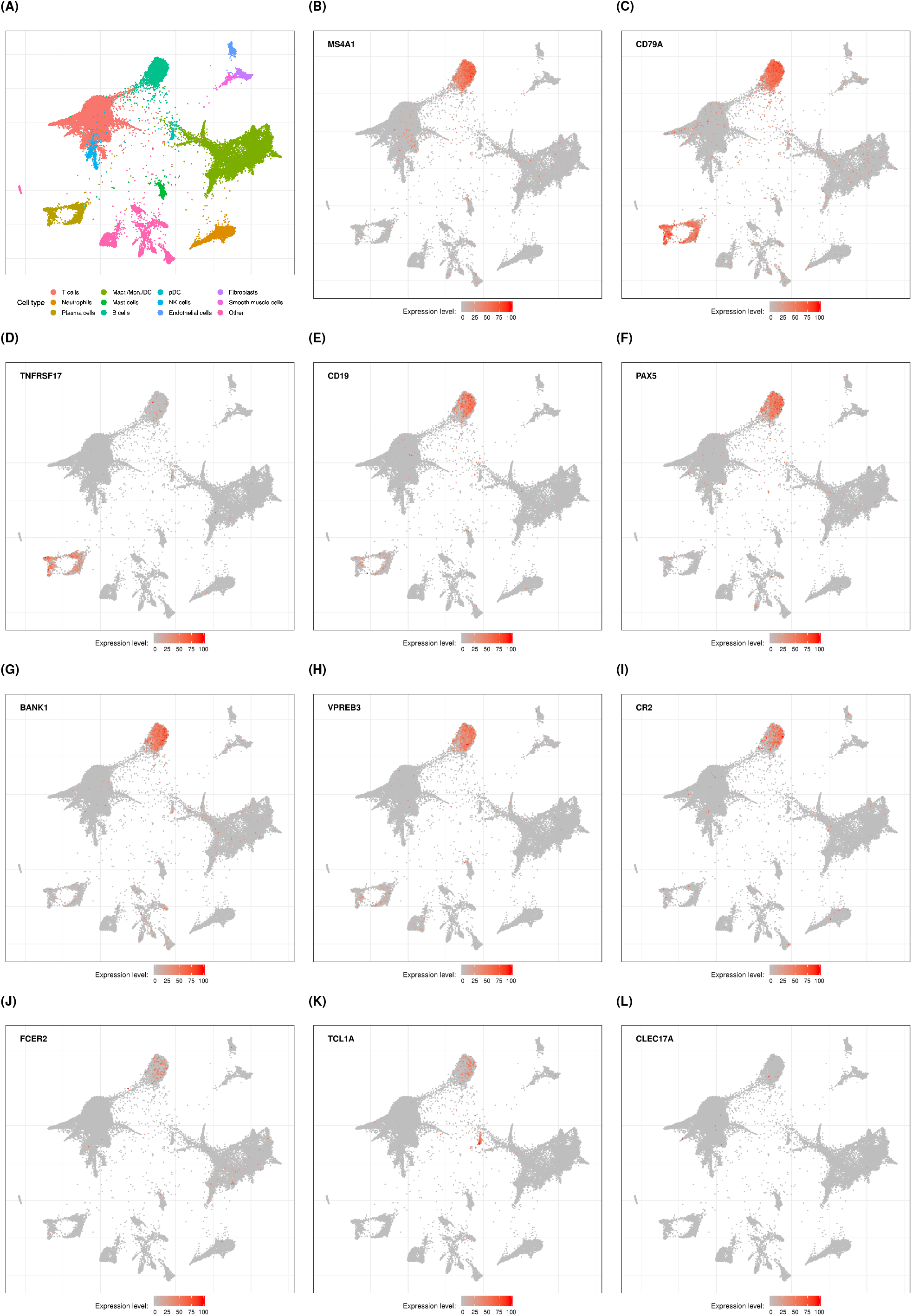
Expression of ICI response associated genes in single cells obtained from lung cancer tumours. Two-dimensional visualizations (SPRING plots) of singlecell RNA sequencing data obtained from lung cancer tumours and published in (25). Each dot represents the transcriptional profile of a single cell. Cells closely associated in each plot are more likely to transcribe similar genes and might thus belong to the same cell type. Expression values and coordinates for each dot were obtained from the original study. The expression level color scale indicates expression level for a particular gene ranging from 0 (minimum expression value observed in the dataset for that gene, usually corresponding to non detectable expression) to 100 (maximum expression value observed in the dataset for that gene). Panels B and C are included in the figure in order to further confirm the identity of cells annotated as “B cells” and “Plasma cells” in panel A. **A.** Cell types annotation as reported in the original study (25). **B.** Expression of *MS4A1/CD20*. **D.** Expression of *CD79A* is detectable in B cells and Plasma cells as expected. **D.** Expression of *TNFRSF17/BCMA* is detectable in Plasma cells but not in B cells as expected. **E.** Expression of *CD19*. **F.** Expression of *PAX5*. **G.** Expression of *BANK1*. **H.** Expression of *VPREB3*. **I.** Expression of *CR2*. **J.** Expression of *FCER2*. **K.** Expression of *TCL1A*, also detectable in Plasmacytoid dendritic cells (pDC, see panel A). **L.** Expression of *CLEC17A*.

**Fig. S8.**
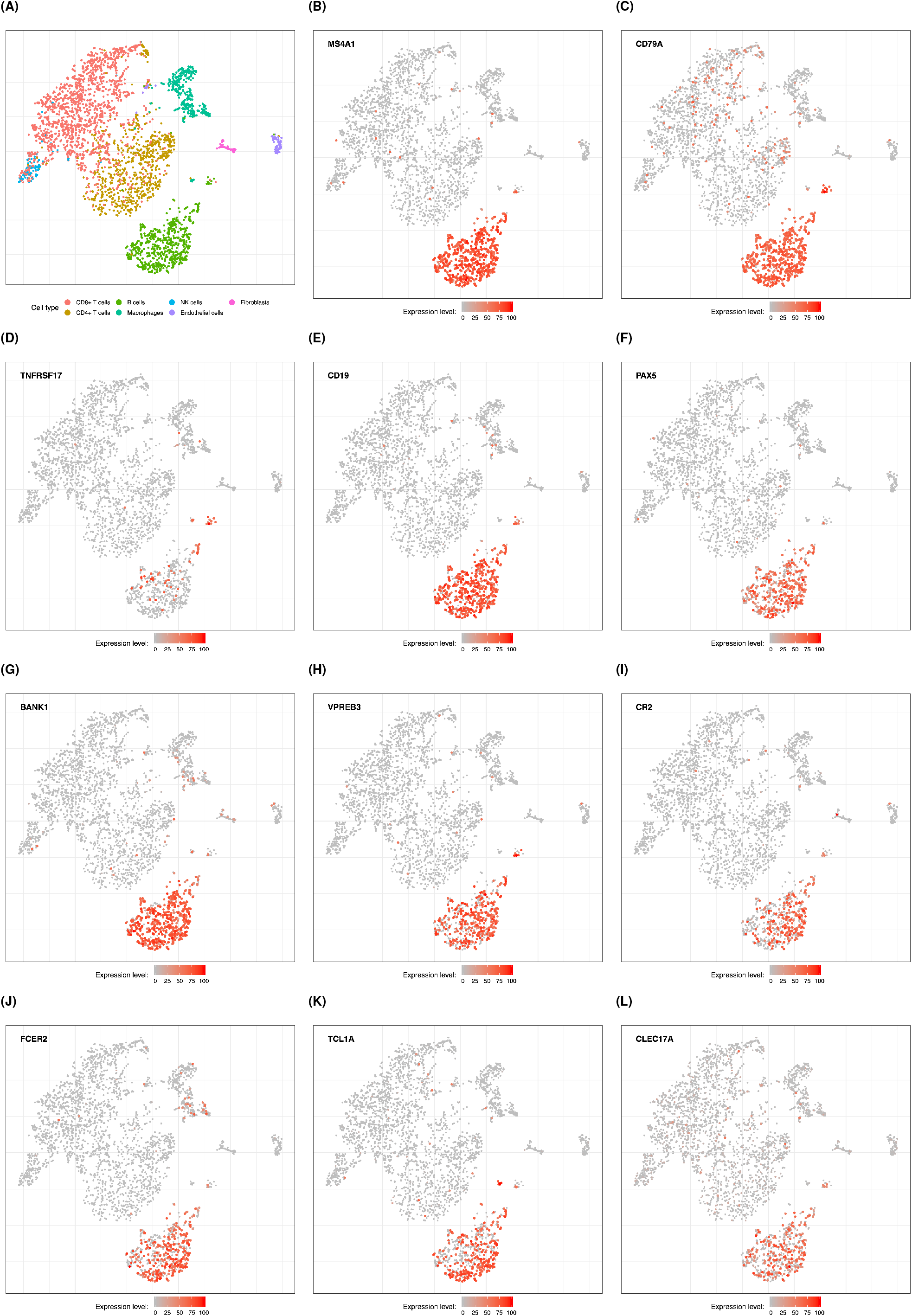
Expression of ICI response associated genes in single cells obtained from melanoma tumours. Two-dimensional visualizations (tSNE plots) of single-cell RNA sequencing data obtained from melanoma tumours and published in (27). Each dot represents the transcriptional profile of a single cell. Cells closely associated in each plot are more likely to transcribe similar genes and might thus belong to the same cell type. Expression values and coordinates for each dot were obtained from the original study. The expression level color scale indicates expression level for a particular gene ranging from 0 (minimum expression value observed in the dataset for that gene, usually corresponding to non detectable expression) to 100 (maximum expression value observed in the dataset for that gene). Panels B and C are included in the figure in order to further confirm the identity of cells annotated as “B cells” and “Plasma cells” in panel A. **A.** Cell types annotation as reported in the original study (27). **B.** Expression of *MS4A1/CD20*. **D.** Expression of *CD79A* **D.** Expression of *TNFRSF17/BCMA*. A small cluster of Plasma cells might be visible but was not clearly annotated in this dataset (see panel A). **E.** Expression of *CD19*. **F.** Expression of *PAX5*. **G.** Expression of *BANK1*. **H.** Expression of *VPREB3*. **I.** Expression of *CR2*. **J.** Expression of *FCER2*. **K.** Expression of *TCL1A*. A small cluster of pDC might be visible but was not clearly annotated in this dataset (see panel A). **L.** Expression of *CLEC17A*.

**Fig. S9.**
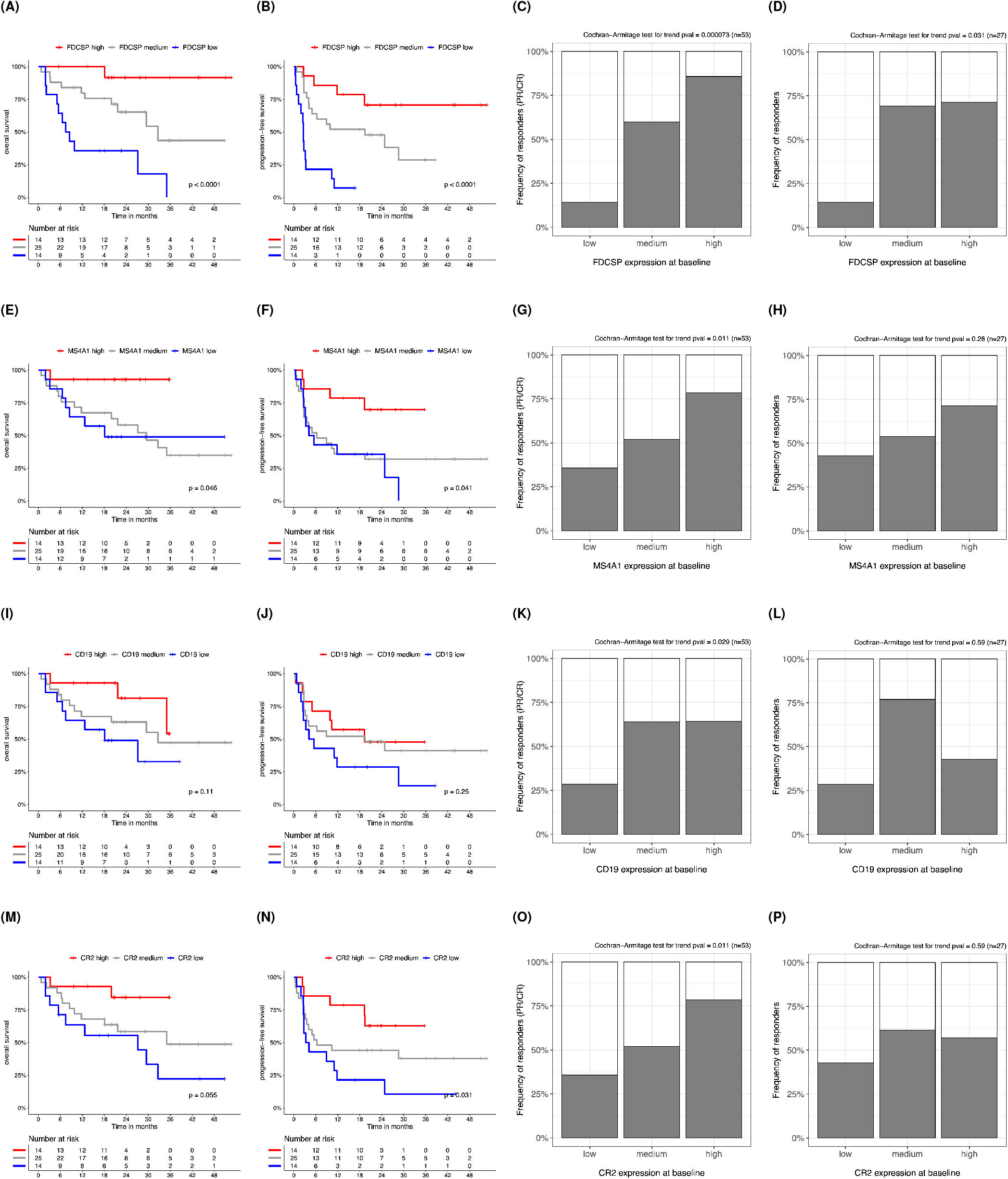
Predicting response in ICI treated patients using pre-treatment expression of FRCs or B cells markers. **A-B.** Kaplan-Meier curves showing different overall survival and progression-free survival in melanoma patients after commencing treatment with ICI. Patients from (10) were stratified in three groups according to pre-treatment expression of *FDCSP*. **C-D.** Association between pre-treatment expression of *FDCSP* and subsequent ICI response in melanoma patients from (10; Panel C) and in clear cell renal cell carcinoma patients from (12; Panel D). Patients were stratified in three groups according to pre-treatment expression of *FDCSP* maintaining the same proportions used in panel A and D. **E-H.** Corresponding results obtained using pre-treatment expression of *MS4A1/CD20*. **I-L.** Corresponding results obtained using pre-treatment expression of *CD19*. **M-P.** Corresponding results obtained using pre-treatment expression of *CR2*.

## Appendix S3: supplementary tables

**Table S2.** Top 50 genes for which pre-treatment expression is associated with subsequent ICI response in melanoma. Genes are ordered by the column *score* which takes into account both p-value and log_2_(*fc*): 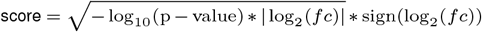. The complete list is provided in a separate spreadsheet file (ICI_resp_baseline.xlsx).

| | gene | $\log_2(f_c)$ | $\log_2(f_c)$ se | p-value | padj | stat | description | score |
| --- | --- | --- | --- | --- | --- | --- | --- | --- |
| 1 | CR2 | 2.5 | 0.27 | 3.3E-21 | 4.9E-17 | 9.5 | complement C3d receptor 2 | 7.1 |
| 2 | FCER2 | 2.2 | 0.32 | 1.3E-13 | 6.3E-10 | 7.4 | Fc fragment of IgE receptor II | 5.3 |
| 3 | CLEC17A | 2 | 0.28 | 1.9E-14 | 1.5E-10 | 7.7 | C-type lectin domain containing 17A | 5.3 |
| 4 | PAX5 | 2.2 | 0.33 | 8.2E-13 | 3.1E-09 | 7.2 | paired box 5 | 5.1 |
| 5 | FDCSP | 2.2 | 0.36 | 6.7E-11 | 1.7E-07 | 6.5 | follicular dendritic cell secreted protein | 4.7 |
| 6 | CD19 | 1.8 | 0.3 | 1.2E-11 | 3.5E-08 | 6.8 | CD19 molecule | 4.5 |
| 7 | TCL1A | 1.9 | 0.34 | 4.2E-10 | 6.3E-07 | 6.2 | T cell leukemia/lymphoma 1A | 4.2 |
| 8 | VPREB3 | 1.7 | 0.31 | 1.7E-10 | 2.9E-07 | 6.4 | V-set pre-B cell surrogate light chain 3 | 4.1 |
| 9 | BANK1 | 1.6 | 0.28 | 1E-10 | 2E-07 | 6.5 | B cell scaffold protein with ankyrin repeats 1 | 4 |
| 10 | MS4A1 | 1.7 | 0.32 | 1.2E-09 | 1.6E-06 | 6.1 | membrane spanning 4-domains A1 | 3.9 |
| 11 | TLR10 | 1.4 | 0.24 | 9E-11 | 1.9E-07 | 6.5 | toll like receptor 10 | 3.8 |
| 12 | BLK | 1.3 | 0.25 | 3.1E-09 | 3.6E-06 | 5.9 | BLK proto-oncogene, Src family tyrosine kinase | 3.3 |
| 13 | MUC7 | 1.7 | 0.44 | 6.6E-07 | 0.00038 | 5 | mucin 7, secreted | 3.3 |
| 14 | PSCA | 1.5 | 0.31 | 4.1E-08 | 3.8E-05 | 5.5 | prostate stem cell antigen | 3.3 |
| 15 | SPIB | 1.3 | 0.26 | 9.9E-09 | 9.9E-06 | 5.7 | Spi-B transcription factor | 3.3 |
| 16 | CYP4F11 | 1.4 | 0.31 | 5.1E-08 | 4.5E-05 | 5.4 | cytochrome P450 family 4 subfamily F member 11 | 3.2 |
| 17 | FAM129C | 1.2 | 0.23 | 5.3E-09 | 5.7E-06 | 5.8 | family with sequence similarity 129 member C | 3.2 |
| 18 | CNR2 | 1.2 | 0.25 | 6.6E-08 | 5.5E-05 | 5.4 | cannabinoid receptor 2 | 2.9 |
| 19 | CUX2 | 1.3 | 0.3 | 3.2E-07 | 0.0002 | 5.1 | cut like homeobox 2 | 2.9 |
| 20 | FCRL1 | 1.3 | 0.33 | 1E-06 | 0.00053 | 4.9 | Fc receptor like 1 | 2.8 |
| 21 | CCR7 | 1.2 | 0.25 | 2E-07 | 0.00014 | 5.2 | C-C motif chemokine receptor 7 | 2.8 |
| 22 | P2RX5 | 1.1 | 0.24 | 2.2E-07 | 0.00015 | 5.2 | purinergic receptor P2X 5 | 2.7 |
| 23 | GPX2 | 1.1 | 0.29 | 3E-06 | 0.0014 | 4.7 | glutathione peroxidase 2 | 2.5 |
| 24 | TSPAN8 | 1.2 | 0.38 | 1.1E-05 | 0.0035 | 4.4 | tetraspanin 8 | 2.5 |
| 25 | TNFRSF13C | 0.98 | 0.22 | 8.7E-07 | 0.00048 | 4.9 | TNF receptor superfamily member 13C | 2.4 |
| 26 | CCL21 | 1.2 | 0.39 | 1.6E-05 | 0.0048 | 4.3 | C-C motif chemokine ligand 21 | 2.4 |
| 27 | ABAT | 0.9 | 0.21 | 1.2E-06 | 0.00061 | 4.9 | 4-aminobutyrate aminotransferase | 2.3 |
| 28 | RGS13 | 1.1 | 0.34 | 1.8E-05 | 0.0051 | 4.3 | regulator of G protein signaling 13 | 2.2 |
| 29 | GP1BA | 0.88 | 0.21 | 2.1E-06 | 0.00096 | 4.7 | glycoprotein Ib platelet subunit alpha | 2.2 |
| 30 | STAP1 | 1 | 0.31 | 1.4E-05 | 0.0044 | 4.3 | signal transducing adaptor family member 1 | 2.2 |
| 31 | CD79A | 1 | 0.37 | 2.7E-05 | 0.0068 | 4.2 | CD79a molecule | 2.2 |
| 32 | SLC27A2 | 1 | 0.32 | 1.8E-05 | 0.0051 | 4.3 | solute carrier family 27 member 2 | 2.2 |
| 33 | C4A | 0.88 | 0.23 | 4.9E-06 | 0.002 | 4.6 | complement C4A (Rodgers blood group) | 2.2 |
| 34 | TREML2 | 0.93 | 0.26 | 9.8E-06 | 0.0033 | 4.4 | triggering receptor expressed on myeloid cells like 2 | 2.2 |
| 35 | APOC4-APOC2 | 1.1 | 0.46 | 5.1E-05 | 0.011 | 4.1 | APOC4-APOC2 readthrough (NMD candidate) | 2.2 |
| 36 | DNASE1L3 | 0.96 | 0.3 | 1.8E-05 | 0.0051 | 4.3 | deoxyribonuclease 1 like 3 | 2.1 |
| 37 | LRMP | 0.78 | 0.2 | 5.5E-06 | 0.0021 | 4.5 | lymphoid restricted membrane protein | 2 |
| 38 | AMDHD1 | 0.83 | 0.25 | 1.7E-05 | 0.0051 | 4.3 | amidohydrolase domain containing 1 | 2 |
| 39 | CPS1 | 0.88 | 0.31 | 3.5E-05 | 0.0085 | 4.1 | carbamoyl-phosphate synthase 1 | 2 |
| 40 | PLAC8 | 0.84 | 0.27 | 2.5E-05 | 0.0066 | 4.2 | placenta associated 8 | 2 |
| 41 | CCR6 | 0.96 | 0.46 | 9.7E-05 | 0.017 | 3.9 | C-C motif chemokine receptor 6 | 2 |
| 42 | SELL | 0.8 | 0.25 | 2E-05 | 0.0054 | 4.3 | selectin L | 1.9 |
| 43 | C4B | 0.78 | 0.25 | 2.5E-05 | 0.0066 | 4.2 | complement C4B (Chido blood group) | 1.9 |
| 44 | MUC3A | 0.77 | 0.27 | 4.1E-05 | 0.0092 | 4.1 | mucin 3A, cell surface associated | 1.8 |
| 45 | VIPR1 | 0.78 | 0.32 | 7.4E-05 | 0.014 | 4 | vasoactive intestinal peptide receptor 1 | 1.8 |
| 46 | IGHV1-58 | 0.89 | 0.62 | 0.00026 | 0.035 | 3.7 | immunoglobulin heavy variable 1-58 | 1.8 |
| 47 | SLC22A1 | 0.79 | 0.37 | 0.00011 | 0.018 | 3.9 | solute carrier family 22 member 1 | 1.8 |
| 48 | CFHR4 | 0.9 | 0.77 | 0.00036 | 0.039 | 3.6 | complement factor H related 4 | 1.8 |
| 49 | ABCG5 | 0.79 | 0.37 | 0.00012 | 0.019 | 3.9 | ATP binding cassette subfamily G member 5 | 1.8 |
| 50 | OIT3 | 0.83 | 0.47 | 0.00018 | 0.027 | 3.7 | oncoprotein induced transcript 3 | 1.8 |

**Table S3.** Multiple linear regression analysis of ICI response, including plasma cell markers. expression of *FDCSP* is significantly associated with ICI response independently of genes expressed in major immune cell types, including Plasma cells. Asterisks indicate the level of statistical significance: *** indicates p-value ≤ 0.001, ** indicates p-value ≤ 0.01, * indicates p-value ≤ 0.05.

|  | Cell type | Markers | Estimate | Std. Error | t value | Pr(>\|t\|) |
| --- | --- | --- | --- | --- | --- | --- |
| 1 | CD8+ T cells | CD8A, CD8B, TRAC | 0.077 | 0.082 | 0.93 | 0.35 |
| 2 | CD4+ T cells | CD4, TRAC | 0.049 | 0.110 | 0.45 | 0.65 |
| 3 | NK cells | GNLY, KLRF1 | -0.074 | 0.092 | -0.80 | 0.42 |
| 4 | B cells | MS4A1, CD19 | 0.078 | 0.053 | 1.50 | 0.14 |
| 5 | Macrophages | CD163, MARCO | -0.019 | 0.049 | -0.40 | 0.69 |
| 6 | Dendritic cells | CD1A, CD1B, CD1E, LILRA4 | -0.059 | 0.075 | -0.78 | 0.44 |
| 7 | Plasma cells | TNFRSF17, MZB1 | -0.093 | 0.056 | -1.70 | 0.098 |
| 8 | FDCSP+ cells | FDCSP | 0.067 | 0.027 | 2.50 | 0.014 * |

**Table S4.** Multiple linear regression analysis of ICI response, excluding *FDCSP*. Expression of B cell markers is significantly associated with ICI response independently of CD8+ T cells and other immune cell types. However, such association becomes statistically non-significant when expression of *FDCSP* is taken into account in the multiple regression model (Table 1 and Table S3). Asterisks indicate the level of statistical significance: *** indicates p-value ≤ 0.001, ** indicates p-value ≤ 0.01, * indicates p-value ≤ 0.05.

|  | Cell type | Markers | Estimate | Std. Error | t value | Pr(>\|t\|) |
| --- | --- | --- | --- | --- | --- | --- |
| 1 | CD8+ T cells | CD8A, CD8B, TRAC | 0.096 | 0.083 | 1.20 | 0.25 |
| 2 | CD4+ T cells | CD4, TRAC | 0.044 | 0.110 | 0.40 | 0.69 |
| 3 | NK cells | GNLY, KLRF1 | -0.045 | 0.093 | -0.48 | 0.63 |
| 4 | B cells | MS4A1, CD19 | 0.120 | 0.051 | 2.20 | 0.027 * |
| 5 | Macrophages | CD163, MARCO | -0.018 | 0.049 | -0.37 | 0.71 |
| 6 | Dendritic cells | CD1A, CD1B, CD1E, LILRA4 | -0.073 | 0.076 | -0.97 | 0.33 |
| 7 | Plasma cells | TNFRSF17, MZB1 | -0.110 | 0.056 | -1.90 | 0.064 |

**Table S5.**
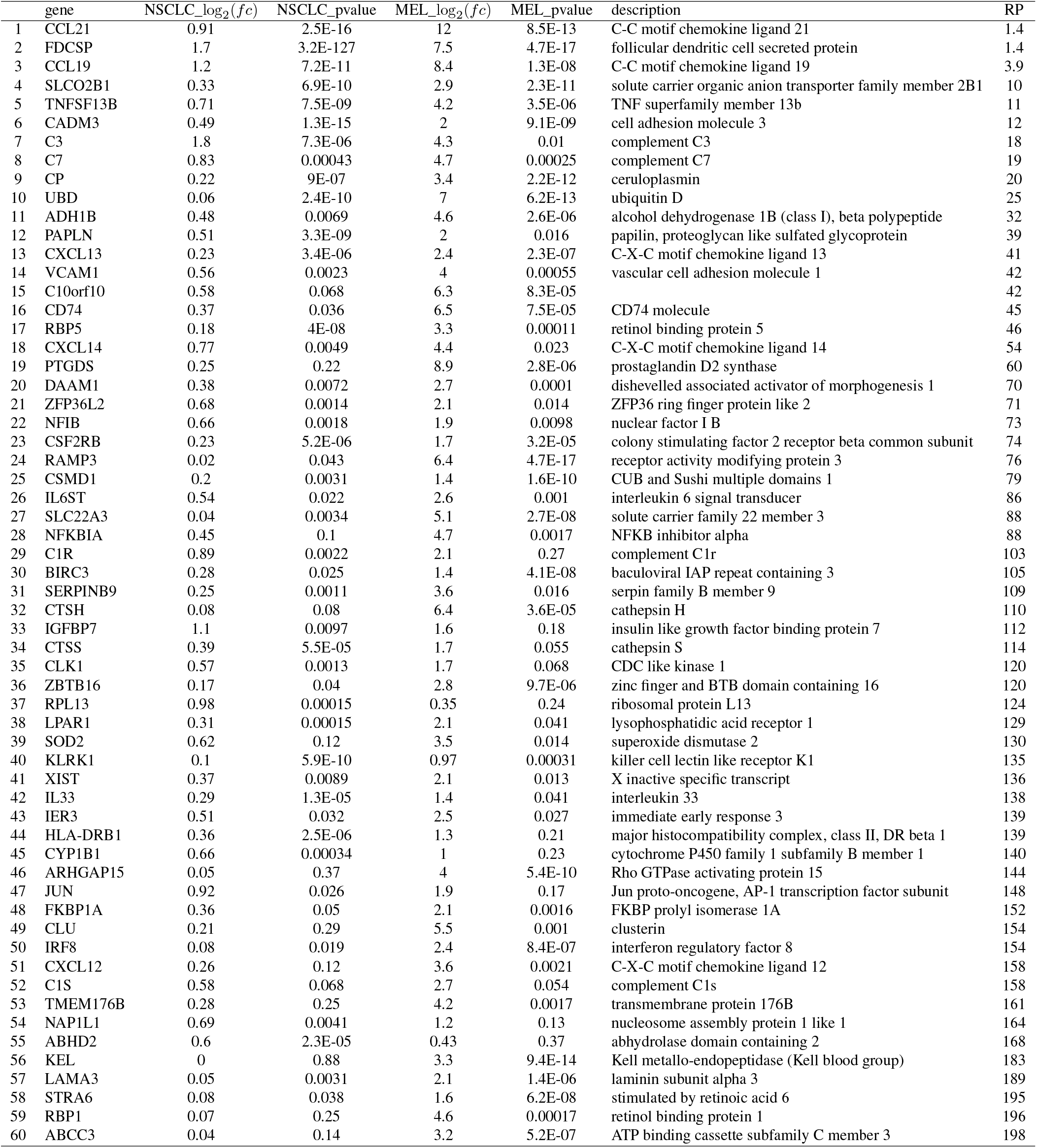
Top 50 genes up-regulated in *FDCSP+* fibroblasts compared to *FDCSP-* fibroblasts detected by using single-cell RNAseq data. Differential gene expression analysis comparing single-cell transcriptomes of *FDCSP+* fibroblasts versus *FDCSP-* fibroblasts obtained from lung cancer (NSCLC) and melanoma tumours (MEL). The analysis was performed using gene expression data published in (25, 27). Results from both datasets were meta-analysed using the RankProduct method (RP; see supplementary methods). The complete list is provided in a separate spreadsheet file (FDCSP_pos_vs_FDCSP_neg_fibroblasts.xlsx).

**Table S6.**
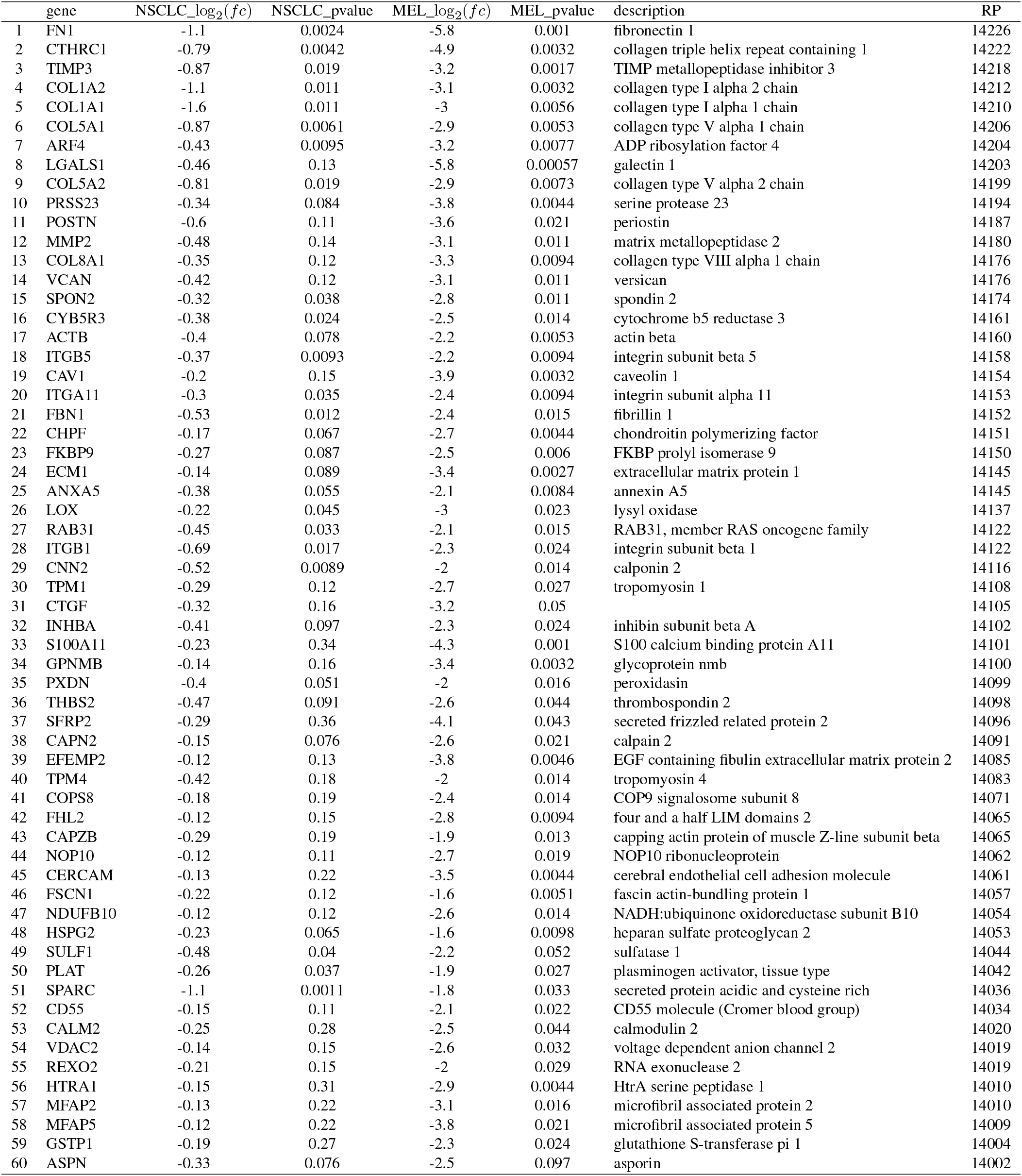
Top 50 genes down-regulated in *FDCSP*+ fibroblasts compared to *FDCSP*- fibroblasts detected by using single-cell RNAseq data. Differential gene expression analysis comparing single-cell transcriptomes of *FDCSP+* fibroblasts versus *FDCSP-* fibroblasts obtained from lung cancer (NSCLC) and melanoma tumours (MEL). The analysis was performed using gene expression data published in (25, 27). Results from both datasets were meta-analysed using the RankProduct method (RP; see supplementary methods). The complete list is provided in a separate spreadsheet file (FDCSP_pos_vs_FDCSP_neg_fibroblasts.xlsx).

**Table S7.**
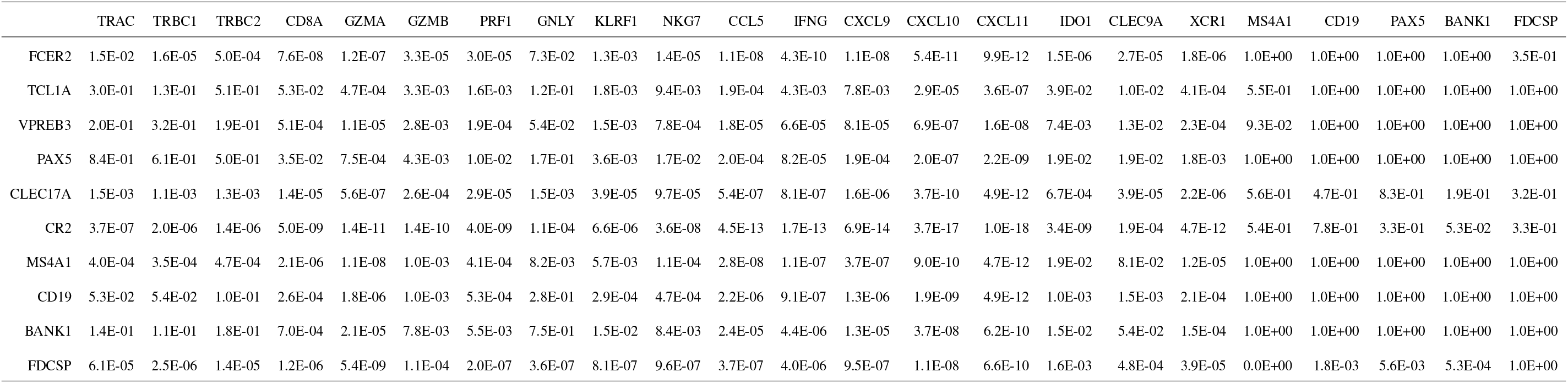
Expression of FDCSP is an independent predictor of response to immune checkpoint inhibitors. FDR values used to generate Fig. 1D

## Bibliography

1. Shalapoνr, S. et al. Immunosuppressive plasma cells impede t-cell-dependent immunogenic chemotherapy. Nature 521, 94–98 (2015).

2. Damsky, W. et al. B cell depletion or absence does not impede anti-tumor activity of pd-1 inhibitors. Journal for immunotherapy of cancer 7, 153 (2019).

3. Hollern, D.P. et al. B cells and t follicular helper cells mediate response to checkpoint inhibitors in high mutation burden mouse models of breast cancer. Cell 179, 1191–1206 (2019).

4. Denton, A. E., Roberts, E. W., Linterman, M. A. & Fearon, D. T. Fibroblastic reticular cells of the lymph node are required for retention of resting but not activated cd8+ t cells. Proceedings of the National Academy of Sciences 111, 12139–12144 (2014).

5. Cremasco, V. et al. B cell homeostasis and follicle confines are governed by fibroblastic reticular cells. Nature immunology 15, 973–981 (2014).

6. Denton, A. E., Carr, E. J., Magiera, L. P., Watts, A. J. & Fearon, D. T. Embryonic fap+ lymphoid tissue organizer cells generate the reticular network of adult lymph nodes. Journal of Experimental Medicine 216, 2242–2252 (2019).

7. Nayar, S. et al. Immunofibroblasts are pivotal drivers of tertiary lymphoid structure formation and local pathology. Proceedings of the National Academy of Sciences 116, 13490–13497 (2019).

8. Helmink, B. A. et al. B cells and tertiary lymphoid structures promote immunotherapy response. Nature (2020).

9. Iglesia, M. D. et al. Genomic analysis of immune cell infiltrates across 11 tumor types. JNCI: Journal of the National Cancer Institute 108 (2016).

10. Ji, R.-R. et al. An immune-active tumor microenvironment favors clinical response to ipili- mumab. Cancer Immunology, Immunotherapy 61, 1019–1031 (2012).

11. Ayers, M. et al. Ifn-γ-related mrna profile predicts clinical response to pd-1 blockade. The Journal of clinical investigation 127, 2930–2940 (2017).

12. Cristescu, R. et al. Pan-tumor genomic biomarkers for pd-1 checkpoint blockade–based immunotherapy. Science 362, eaar3593 (2018).

13. Cabrita, R. et al. Tertiary lymphoid structures improve immunotherapy and survival in melanoma. Nature (2020).

14. Petitprez, F. et al. B cells are associated with survival and immunotherapy response in sarcoma. Nature (2020).

15. Schwartz, L. H. et al. Recist 1.1—update and clarification: From the recist committee. European journal of cancer 62, 132–137 (2016).

16. Kassambara, A. et al. Genomicscape: an easy-to-use web tool for gene expression data analysis. application to investigate the molecular events in the differentiation of b cells into plasma cells. PLoS computational biology 11, e1004077 (2015).

17. Marshall, A. J. et al. Fdc-sp, a novel secreted protein expressed by follicular dendritic cells. The Journal of Immunology 169, 2381–2389 (2002).

18. Al-Alwan, M. et al. Follicular dendritic cell secreted protein (fdc-sp) regulates germinal center and antibody responses. The Journal of Immunology 178, 7859–7867 (2007).

19. Zilionis, R. et al. Single-cell transcriptomics of human and mouse lung cancers reveals conserved myeloid populations across individuals and species. Immunity 50, 1317–1334 (2019).

20. Jerby-Arnon, L. et al. A cancer cell program promotes t cell exclusion and resistance to checkpoint blockade. Cell 175, 984–997 (2018).

21. Hartung, E. et al. Induction of potent CD8 t cell cytotoxicity by specific targeting of antigen to cross-presenting dendritic cells in vivo via murine or human XCR1. The Journal of Immunology 194, 1069–1079 (2014).

22. Chow, M. T. et al. Intratumoral activity of the cxcr3 chemokine system is required for the efficacy of anti-pd-1 therapy. Immunity 50, 1498–1512 (2019).

23. Graham, S. A. et al. Prolectin, a glycan-binding receptor on dividing b cells in germinal centers. Journal of Biological Chemistry 284, 18537–18544 (2009).

24. Takahashi, K. et al. Mouse complement receptors type 1 (cr1; cd35) and type 2 (cr2; cd21): expression on normal b cell subpopulations and decreased levels during the development of autoimmunity in mrl/lpr mice. The Journal of Immunology 159, 1557–1569 (1997).

25. Gide, T. N. et al. Distinct immune cell populations define response to anti-pd-1 monotherapy and anti-pd-1/anti-ctla-4 combined therapy. Cancer cell 35, 238–255 (2019).

26. Miao, D. et al. Genomic correlates of response to immune checkpoint therapies in clear cell renal cell carcinoma. Science 359, 801–806 (2018).

27. Garnelo, M. et al. Interaction between tumour-infiltrating b cells and t cells controls the progression of hepatocellular carcinoma. Gut 66, 342–351 (2017).

28. Gunn, M. D. et al. Mice lacking expression of secondary lymphoid organ chemokine have defects in lymphocyte homing and dendritic cell localization. The Journal of experimental medicine 189, 451–460 (1999).

29. Förster, R. et al. Ccr7 coordinates the primary immune response by establishing functional microenvironments in secondary lymphoid organs. Cell 99, 23–33 (1999).

## Supplementary methods references

1. Akbani, R. et al. Genomic classification of cutaneous melanoma. Cell 161, 1681–1696 (2015).

2. Grossman, R. L. et al. Toward a shared vision for cancer genomic data. New England Journal of Medicine 375, 1109–1112 (2016).

3. Mailman, M. D. et al. The ncbi dbgap database of genotypes and phenotypes. Nature genetics 39, 1181–1186 (2007).

4. Van Allen, E. M. et al. Genomic correlates of response to ctla-4 blockade in metastatic melanoma. Science 350, 207–211 (2015).

5. Edgar, R., Domrachev, M. & Lash, A. E. Gene expression omnibus: Ncbi gene expression and hybridization array data repository. Nucleic acids research 30, 207–210 (2002).

6. Riaz, N. et al. Tumor and microenvironment evolution during immunotherapy with nivolumab. Cell 171, 934–949 (2017).

7. Hugo, W. et al. Genomic and transcriptomic features of response to anti-pd-1 therapy in metastatic melanoma. Cell 165, 35–44 (2016).

8. Auslander, N. et al. Robust prediction of response to immune checkpoint blockade therapy in metastatic melanoma. Nature medicine 24, 1545–1549 (2018).

9. Leinonen, R. et al. The european nucleotide archive. Nucleic acids research 39, D28–D31 (2010).

10. Gide, T. N. et al. Distinct immune cell populations define response to anti-pd-1 monotherapy and anti-pd-1/anti-ctla-4 combined therapy. Cancer cell 35, 238–255 (2019).

11. Helmink, B. A. et al. B cells and tertiary lymphoid structures promote immunotherapy response. Nature (2020).

12. Miao, D. et al. Genomic correlates of response to immune checkpoint therapies in clear cell renal cell carcinoma. Science 359, 801–806 (2018).

13. Cunningham, F. et al. Ensembl 2019. Nucleic Acids Research 47, D745–D751 (2018). URL https://doi.org/10.1093/nar/gky1113.

14. Bray, N. L., Pimentel, H., Melsted, P. & Pachter, L. Near-optimal probabilistic RNA-seq quantification. Nature Biotechnology 34, 525–527 (2016). URL https://doi.org/10.1038/nbt.3519.

15. Love, M. I., Huber, W. & Anders, S. Moderated estimation of fold change and dispersion for RNA-seq data with DESeq2. Genome Biology 15 (2014). URL https://doi.org/10.1186/s13059-014-0550-8.

16. Soneson, C., Love, M. I. & Robinson, M. D. Differential analyses for RNA-seq: transcript-level estimates improve gene-level inferences. F1000Research 4, 1521 (2015). URL https://doi.org/10.12688/f1000research.7563.1.

17. Love, M. I., Anders, S. & Huber, W. Analyzing rna-seq data with deseq2 (2019). URL http://bioconductor.org/packages/devel/bioc/vignettes/DESeq2/inst/doc/DESeq2.html#tximport.

18. R Core Team. R: A Language and Environment for Statistical Computing. R Foundation for Statistical Computing, Vienna, Austria (2018). URL https://www.R-project.org/.

19. Huber, W., von Heydebreck, A., Sültmann, H., Poustka, A. & Vingron, M. Parameter estimation for the calibration and variance stabilization of microarray data. Statistical applications in genetics and molecular biology 2 (2003).

20. Johnson, W. E., Li, C. & Rabinovic, A. Adjusting batch effects in microarray expression data using empirical bayes methods. Biostatistics 8, 118–127 (2007).

21. Leek, J. T. et al. Sva: surrogate variable analysis. 2015. R package version 3, 25–27.

22. Schwartz, L. H. et al. Recist 1.1—update and clarification: From the recist committee. European journal of cancer 62, 132–137 (2016).

23. Kassambara, A., Kosinski, M., Biecek, P. & Fabian, S. Package ‘survminer’. Drawing Survival Curves using ‘ggplot2’.(R package version 0.3. 1.) (2017).

24. Peto, R. & Peto, J. Asymptotically efficient rank invariant test procedures. Journal of the Royal Statistical Society: Series A (General) 135, 185–198 (1972).

25. Zilionis, R. et al. Single-cell transcriptomics of human and mouse lung cancers reveals conserved myeloid populations across individuals and species. Immunity 50, 1317–1334 (2019).

26. Weinreb, C., Wolock, S. & Klein, A. M. Spring: a kinetic interface for visualizing high dimensional single-cell expression data. Bioinformatics 34, 1246–1248 (2018).

27. Jerby-Arnon, L. et al. A cancer cell program promotes t cell exclusion and resistance to checkpoint blockade. Cell 175, 984–997 (2018).

28. Maaten, L. v. d. & Hinton, G. Visualizing data using t-sne. Journal of machine learning research 9, 2579–2605 (2008).

29. Mann, H. B. & Whitney, D. R. On a test of whether one of two random variables is stochastically larger than the other. The annals of mathematical statistics 50–60 (1947).

30. Breitling, R., Armengaud, P., Amtmann, A. & Herzyk, P. Rank products: a simple, yet powerful, new method to detect differentially regulated genes in replicated microarray experiments. FEBS letters 573, 83–92 (2004).

31. Hong, F. & Breitling, R. A comparison of meta-analysis methods for detecting differentially expressed genes in microarray experiments. Bioinformatics 24, 374–382 (2008).

